# Retinal microglia-derived S100A9 incite NLRP3 inflammasome in a Western diet fed Ossabaw pig retina

**DOI:** 10.1101/2024.10.30.621160

**Authors:** Rayne R Lim, Anju Thomas, Aparna Ramasubramanian, Shyam S. Chaurasia

## Abstract

**Purpose:** We established S100A9 as a myeloid-derived damage-associated molecular pattern (DAMPs) protein associated with increasing severity of diabetic retinopathy (DR) in type 2 diabetic subjects. The present study investigates the retinal localization, expression, and mechanisms of action for S100A9 in the young obese Ossabaw pig retina.

**Methods:** Retinae from Ossabaw pigs fed a Western diet for 10 weeks were evaluated for S100 and inflammatory mediator expression using quantitative PCR and Western blot. Double immunohistochemistry was performed to identify the cellular sources of S100A9 in the pig retina. Primary pig retinal microglial cells (pMicroglia) were examined for S100A9 production. S100A9-induced responses were also investigated, and inhibitor studies elucidated the mechanism of action via the NLRP3 inflammasome. A specific inhibitor, Paquinimod (ABR-215757), was administered *in vitro* to assess the rescue of S100A9-induced NLRP3 inflammasome activation in pMicroglia.

**Results:** The expression of the S100 family in the obese Ossabaw pig retina showed a significant elevation of S100A9, consistent with increased levels of circulating S100A9. Moreover, the retina had elevated levels of inflammatory mediators IL-6, IL-8, MCP-1, IL-1β and NLRP3. Retinal microglia in obese Ossabaw were activated and accompanied by an increased expression of intracellular S100A9. pMicroglia isolated from pig retina transformed from ramified to amoeboid state when activated with LPS and produced high S100A9 transcript and protein levels. The S100A9 protein, in turn, further activated pMicroglia by heightened production of S100A9 transcripts and secretion of pro-inflammatory IL-1β protein. Inhibition of TLR4 with TAK242 and NLRP3 with MCC950 attenuated the production of IL-1β during S100A9 stimulus. Finally, pre-treatment with Paquinimod successfully reduced S100A9-driven increases of glycosylated-TLR4, NLRP3, ASC, Caspase-1, and IL-1β production.

**Conclusion:** We demonstrated that microglial-derived S100A9 perpetuates pro-inflammatory responses via the NLRP3 inflammasome in the retina of young Western-diet-fed Ossabaw pigs exhibiting diabetic retinopathy.

## Introduction

Diabetic retinopathy (DR) manifests in varying stages of severity in approximately one-third of the diabetic population^1^, making it one of the leading causes of blindness in working adults^2^. The retinal disease embarks in the non-proliferative stage (NPDR) and gradually deteriorates to proliferative DR (PDR). Current treatment options target the late stages of the disease progression via intraocular anti-VEGF treatments and pan-retinal laser photocoagulation. Newer therapies targeting other molecular pathways are unmet needs in the early stages of DR, including the inflammatory milieu produced during NPDR^3^ and the non-responsive patient population to anti-VEGF treatments^4,5^.

Systemic uncontrolled hyperglycemia resulting in chronic sterile inflammation is a critical pathological component in DR progression. Pro-inflammatory cytokines, including IL-1β, TNFα^6,7^, and chemokines, are a biochemical hallmark in humans^8–11^ and animal models^12,13^. While corticosteroids were an effective alternative in anti-VEGF resistance patients^4,5,14^, therapeutic inhibition of the inflammatory pathway in humans using NSAIDS^15^ or anti-TNFα^16^ was met with limited success and conflicting results in diabetic macular edema. Identifying inflammatory initiators describes the damage-associated molecular patterns (DAMPs), endogenous molecules released during cell stress that exacerbate inflammation by instigating a damage chain reaction^17^. Members of this family include high-mobility group box 1 (HMGB1)^18^ and the S100 proteins^19^, which were found elevated in the human diabetic retina.

S100A9 is a cytosolic Ca^2+^-binding protein constitutively expressed by neutrophils and monocytes^20^. It involves several cellular functions, including proliferation, migration, chemotaxis, and, most notably, inflammatory pathways^21^. S100A9 is an early indicator for several inflammatory-associated diseases, as it is produced by cancer^22^ and epithelial^21^ cells during the inflammatory phase of diseases and is accumulated in locally afflicted tissue niches^24^. S100A9 is hence reported to be associated with inflammatory bowel disease^25^, rheumatic diseases^26^, lupus nephritis^27^, and diabetes^28,29^. We recently described increased systemic levels of S100A9 with the progression of DR in human T2DM patients^30^, while protein expression has also been documented in the vitreous and epiretinal membranes of PDR patients^19^. However, the retinal cellular source and mechanism of action of S100A9 is yet to be determined. Moreover, no strategies have yet been uncovered for attenuating DAMPs activity in the diabetic retina.

Microglia are the only known myeloid-derived cells in the retina. They retain functions as resident macrophages and perform immune surveillance due to their motility and proximity to the cells across the inner retina^31^. Microglia activation has been described as an early event in the pathogenesis of DR, and their increased cell numbers and aggravation around microvascular abnormalities^32^ are suggested to be potential prognosis markers^33^. Oral administration of minocycline has been found to attenuate activation of microglial cells and improve visual acuity in patients with diabetic macular edema (DME)^34^, suggesting a possible attenuation of the exacerbated inflammatory response in the retina via microglia regulatory strategies. Moreover, a recent transcriptome study of freshly isolated CD11b+ CD45^lo^ brain microglial cells found S100A9 to be selectively expressed in these cells^35^. The microglial transformation from a ramified to amoeboid morphology in the activated stage during DR produces pro-inflammatory cytokines, including IL-1β^31^, contributing to the disease progression in the retina. We recently described microglia-induced secretion of IL-1β via the activation of the nod-like receptor (NLR), a cytosolic DAMPs receptor in mouse models of PDR^36^, suggesting their involvement in the progression of the disease.

The role of microglia in the immune system is primarily studied in rodents, limiting our understanding of their immune cell markers and regulation in humans, especially in the host defense ligand and receptor family^37^. The pig immune system resembles humans more than rodents^37,38^ due to genomic similarities to humans and mice^39^. The pig retina also resembles humans in ultrastructure^40^, vasculature^41^, and topography^42^. Of note, the pig has a pseudo-macular region (absent in rodents) that is devoid of the major blood vessel and contains a high density of ganglion cells and visual cones^43^, which places it in a unique position between rodents and non-human primates, especially for pharmacological and preclinical trials. Recent reports have outlined pig models in ophthalmology research for continuous tracking of common pathological changes due to hyperglycemia, dyslipidemia, inflammation, and oxidative stress in the neuronal cells and vasculature of the retina^44,45^. Recently, we characterized a novel *in vivo* model of diabetic retinopathy (DR) in young Ossabaw pigs fed a western diet, which showed metabolic syndrome and multiple early neurodegenerative features of DR seen in human patients^46^.

This study was designed to examine S100A9 in the diabetic Ossabaw pig retina, evaluate microglial-derived S100A9 in an *in vitro* microglial culture, and investigate the inhibition of S100A9 using Paquinimod, a specific inhibitor, to understand the contribution of S100A9 to the inflammatory pathology in DR. We hypothesize that activated retinal microglial cells produce S100A9 under cellular stress to exert pro-inflammatory effects on neighboring microglia cells, which can be inhibited by Paquinimod via downregulation of NLRP3 inflammasome, thus attenuating chronic retinal inflammation during DR pathogenesis.

## Material and Methods

### Animals

Ossabaw pigs were divided into two groups at 3.5 months of age into (1) Lean and (2) Obese groups, as previously described^46^. Obese pigs were fed 10 weeks of a “Western” diet (5B4L, Lab Diet; 4.14 kcal/g, 40.8% from carbohydrates, 16.2% from protein, and 43% from fat) as described^44^. Pigs were euthanized at 6 months of age, and eyes were enucleated for immunohistochemistry.

Domestic pigs (American Landrace) for microglia culture were sourced from the local abattoir at the University of Missouri, Columbia, MO. Previous reports did not find significant differences in monocyte markers or function across breeds of pigs^47^. Pigs were euthanized humanely, and eyes were enucleated within 30 minutes of euthanasia.

### Immunohistochemistry

Ossabaw pig eyes were fixed in 4% PFA and embedded in OCT. Twelve-micron thick retinal sections were collected, and immunostaining was performed using a standard protocol. The blocking buffer is comprised of 5% donkey serum. Antibodies used were Iba-1 (1:500; Wako Pure Chemical Industries, Osaka, Japan) and S100A9 (1:50; Novus Biologicals, Littleton, CO, USA) as described in Table 1.

**Table 1.** Antibodies and dilutions for immunostaining and western blot.

| Target | Dilution | Catalogue Number | Company |
| --- | --- | --- | --- |
| **Immunostaining** |  |  |  |
| S100A9 | 1:50 | NB110-89726 | Novus Biologicals |
| Iba-1 | 1:100 | 019-19741 | Wako |
| **Western blot** |  |  |  |
| S100A9 | 1:200 | - | Dr Phillipe Tessier (Université Laval, Québec, Canada) |
| NLRP3 | 1:500 | 15101 | Cell Signaling Technology |
| ASC | 1:500 | AG-25B-0006 | Adipogen |
| Caspase-1 | 1:200 | AG-20B-0042 | Adipogen |
| TLR4 | 1:50 | sc-10741 | Santa Cruz |
| IL-1β | 1:50 | MP425 | Thermo Fisher Scientific |
| β-actin | 1:2000 | 3700 | Cell Signaling Technology |
| GAPDH | 1:5000 | sc-25778 | Santa Cruz |

### Cell culture

Pig retinal microglia (pMicroglia) were cultured as described^48^. In brief, pig eyes were hemisected, and the retina was isolated under aseptic conditions. Retina tissue was triturated, digested with 0.2% collagenase A (Roche, Indianapolis, IN, USA), strained, and filtrate cultured in DMEM-HG with 10% FBS, 1% pen/strep, gentamicin (25ng/ml), amphotericin B (10ng/ml) and recombinant human (rh) M-CSF (25ng/ml, Peprotech, Rocky Hill, NJ, USA) at 3-4×10^5^ cells/cm^2^ on T75 flasks. To isolate pMicroglia, flasks were shaken between days 16 to 21 for 2 hours on an orbital shaker in the incubator. The cell culture supernatant was collected, spun down, and seeded at 5×10^4^ cells/cm^2^ for subsequent experiments.

### pMicroglia treatment

pMicroglia were seeded on 24-well plates for real-time PCR/ELISA and 60mm dishes for Western blot assays. On day 2, cells were treated with serum-free DMEM-HG containing no phenol red and 1X GlutaMAX for 24 hours with LPS (Invivogen, San Diego, CA, USA) dosage ranging from 0.01ng/ml to 100ng/ml, or S100A9 (ProSpec-Tany TechnoGene Ltd, Rehovot, Israel) doses ranging from 0.01nM to 1000nM. ATP (5mM, Invivogen) was given 30 minutes prior to the collection of cells. For the mechanistic studies, cells were pretreated with inhibitors for 1 hour prior to the addition of S100A9 (100nM). The experiment was terminated after 24 hours, and cells were collected for transcript or protein evaluation. Inhibitors used were TAK-242 (1μM; Calbiochem, MilliporeSigma, Burlington, MA, USA), MCC950 (1μM; Calbiochem), and Paquinimod (ABR-215757; Active Biotech AB, Lund, Sweden). ATP (5mM) was given 15 minutes before collection.

### Real-time PCR (qPCR)

pMicroglia were collected in RLT buffer with 1% β-mercaptoethanol, lysed using a QIA shredder spin column, and RNA isolated using the RNeasy mini kit. Reverse transcription was performed with 250ng RNA using GoScript (Promega, Madison, WI, USA), and qPCR was performed in a 384-well plate on the ViiA7 (Applied Biosystems, Foster City, CA, USA) using PowerUp SYBR green dye (Thermo Fisher Scientific). Pig and mouse primer sequences are described in Table 2.

**Table 2.**
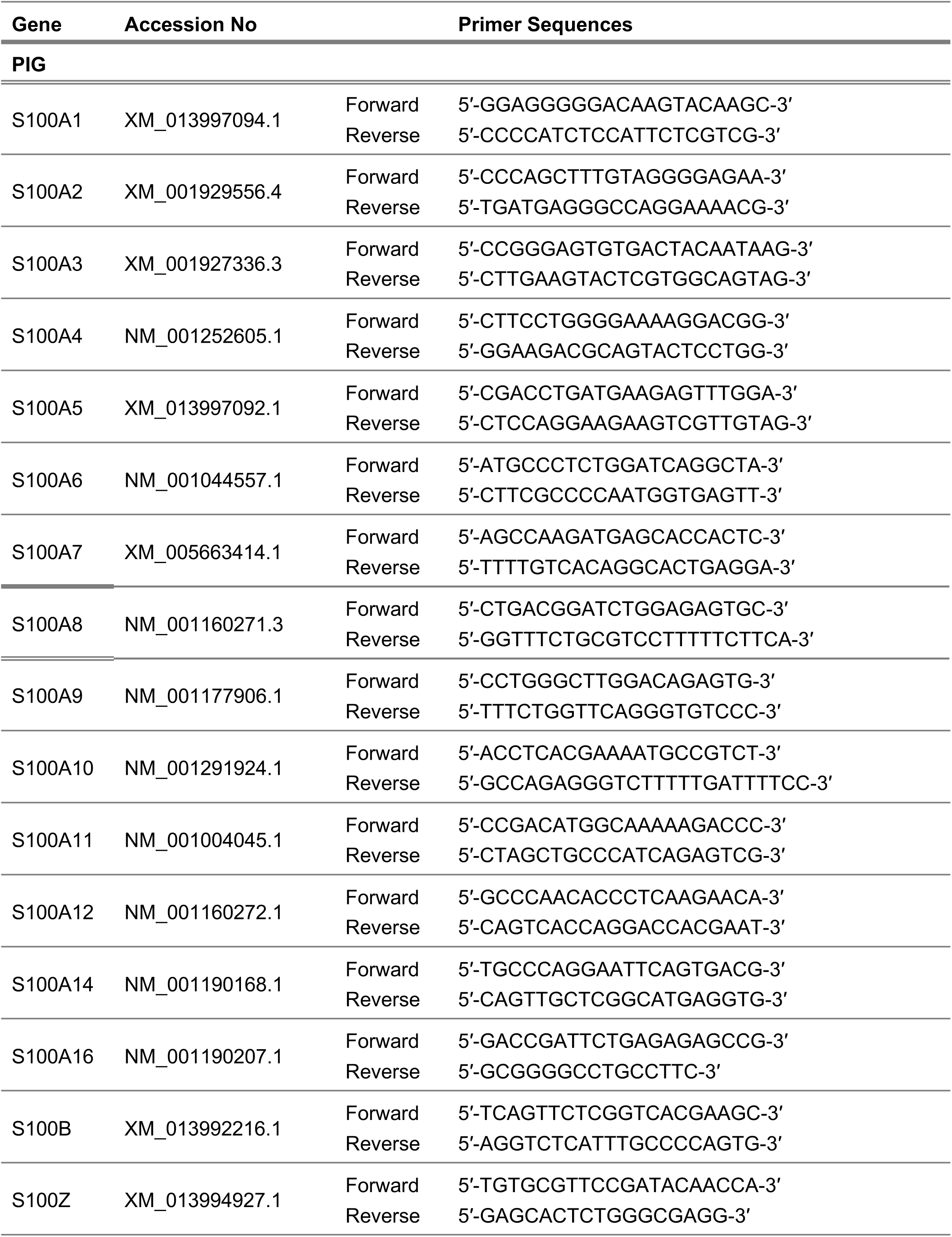
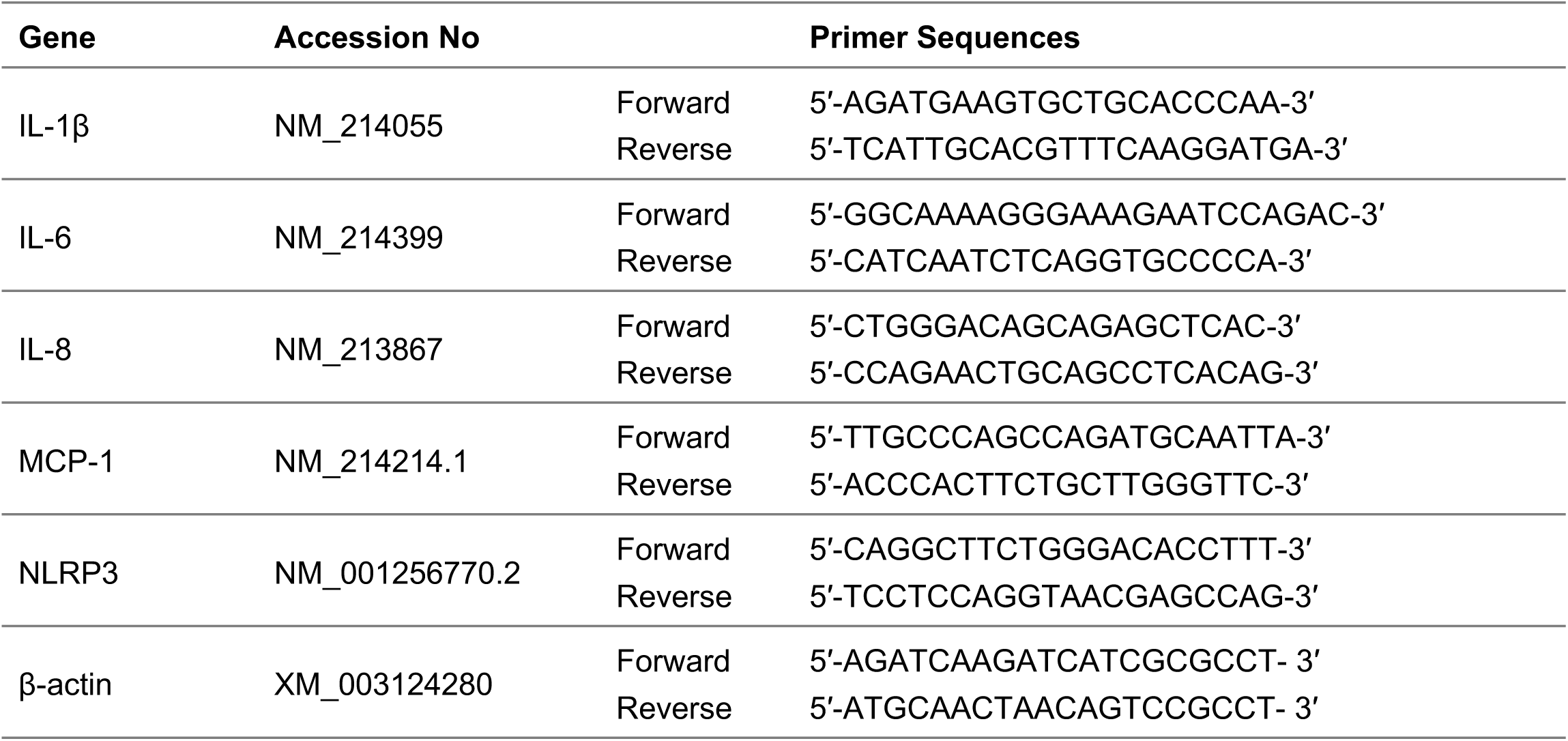
Real-time PCR primer sequences.

### Western blot

Retinal tissues (10mg) homogenized in 200µl RIPA buffer with Halt protease inhibitors were spun down to remove cell debris. Cells from culture plates were lysed in RIPA buffer and collected with a rubber scraper. Cell protein lysate was sonicated for 3 x 15sec at 20% amplitude and spun down to remove cell debris. Quantification was performed using the BCA protein assay kit (Thermo Fisher Scientific). 50µg of cell lysate was run on a 12% gel and transferred onto a nitrocellulose membrane. Blocking of blots was done with 5% milk in 1X TBST. The primary antibody was incubated overnight with shaking at 4°C, while the secondary antibody was added at room temperature for 1 hour. Antibody dilutions are shown in Table 1. Blots were detected using chemiluminescence on the C-DiGit system (Li-Cor, Lincoln, NE, USA).

### Enzyme-linked immunoassay (ELISA)

To quantify cytokines secreted in the media, ATP (5mM) was added 30 minutes before collection. The media was spun down at 10,000 rpm at 4°C for 10 minutes to remove cellular debris and stored at –80°C for further analysis. Porcine IL-1β, IL-6, IL-8, and Mouse IL-1β ELISAs (R&D Systems, Minneapolis, MN, USA) were performed according to the manufacturer’s instructions. 100µl of cell culture media was used in duplicate.

### Cell Viability assay

pMicroglia were seeded in a 96-well plate and allowed to adhere for 24 hours. They were then given paquinimod in varying concentrations (10 – 1000µM). Cell viability was measured after 24 hours using PrestoBlue Cell Viability Reagent (Thermo Fisher Scientific). Treatment media was replaced with 1X PrestoBlue reagent diluted in phenol red-free media. After 2 hours of incubation, absorbance was measured at 570nm, with reference values at 600nm. The experiment was repeated twice in duplicates.

### Statistical analysis

Data were analyzed and plotted using GraphPad software (GraphPad Prism 6.0; GraphPad Software, Inc., La Jolla, CA, USA) and represented as mean±SEM. Student’s *t*-tests were performed to compare the two groups; one-way ANOVA with post-hoc Bonferonni correction was performed for comparisons with more than two groups. P<0.05 was considered statistically significant.

## Results

### Increased inflammatory factors and S100A9 in obese Ossabaw pigs

The S100 family has sixteen members, each serving diverse roles beyond regulating inflammation. Transcript expression revealed expression of all S100 family members in the Ossabaw pig retina (Fig. 1A). Surprisingly, S100A9 alone was highly upregulated in the obese pig retina. Additionally, ELISA of plasma S100A9 revealed a notable increase in circulating plasma levels (Fig. 1B), similar to previous human clinical study^30^. Staining of the pig retinas found S100A9 localization predominantly in the ganglion cell layer (GCL) and nerve fiber layer (NFL) but was increased across the retina and the RPE in the obese pig (Fig. 1C).

**Figure 1.**
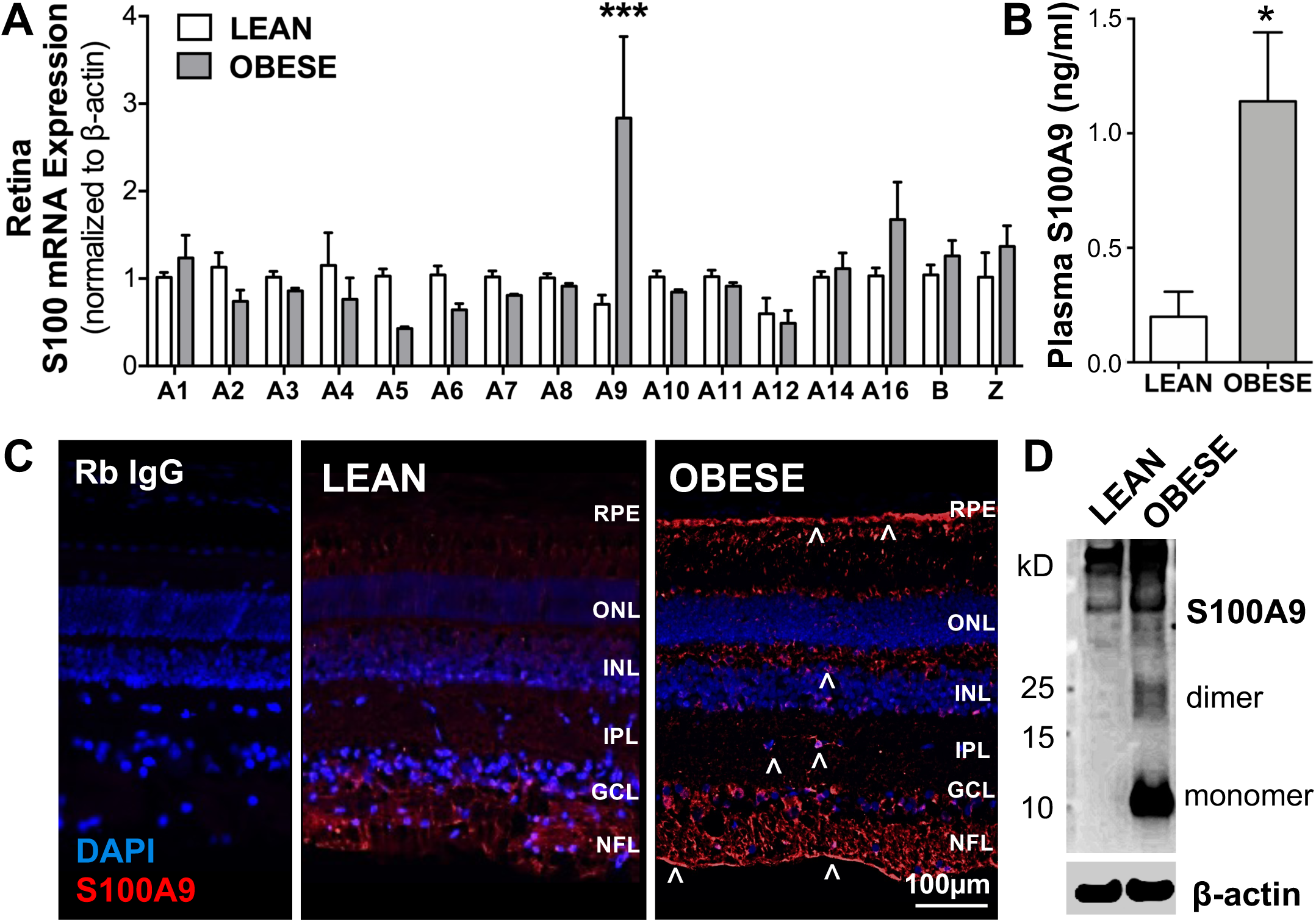
S100A9 is upregulated in the retina of Ossabaw pigs fed a Western diet. (**A**) Real-time PCR of retinal S100 profile in lean and obese Ossabaw pigs showed significant upregulation of S100A9 in pigs fed with Western diet. (**B**) Plasma samples showed elevated levels of S100A9 in obese pigs. (**C)** Immunostaining showed S100A9 increase to be localized to the retinal pigmented epithelium (RPE), inner and outer plexiform layer (IPL & OPL) of the obese pigs. n = 3 eyes per group. *, P<0.05 & ***, P<0.001 vs. Lean. (**D**) Retinal lysate ran on western blot similar had a large increase in S100A9 protein, seen as a monomeric band at 13kD and a dimeric band at 26kD.

S100A9 dimerizes target binding but also forms tetramers and high-order complexes due to their high intrinsic amyloid-forming capacity^49^. S100A9 band consistent with the multimeric size was seen in both the lean and obese pig retina. However, significantly elevated monomeric and dimeric S100A9 levels were seen in the obese Ossabaw pig retina (Fig. 1D).

Since inflammation is a critical factor in DR pathogenesis, the retinas of lean and obese animals were evaluated for factors commonly associated with diabetic retinas. Real-time PCR showed increased production of IL-6, IL-8, and MCP-1 transcripts (Supp Fig. 1A). IL-1β and NLRP3 were upregulated (Supp Fig. 1B).

### Ossabaw pigs fed a western diet expressed high levels of S100A9 in the retinal microglial cells

To identify the cells responsible for S100A9 localization in the retina, double immunohistochemistry was performed with Iba-1, a classical microglial marker. Microglia activation was seen in the obese pig retina compared to the lean pig retina, with changes in cell morphology and increased numbers in the outer plexiform layer (OPL) (Fig. 2A-F). Though S100A9 was seen distributed across the inner retina, the protein was distinctly identified in the cell body of ramified microglia in the lean pig retina (Fig. 2C). Expression was subsequently increased in the amoeboid microglia of the diabetic retina (Fig. 2F). Microglia in the OPL of obese animals were also found to express S100A9 with greater intensity.

**Figure 2.**
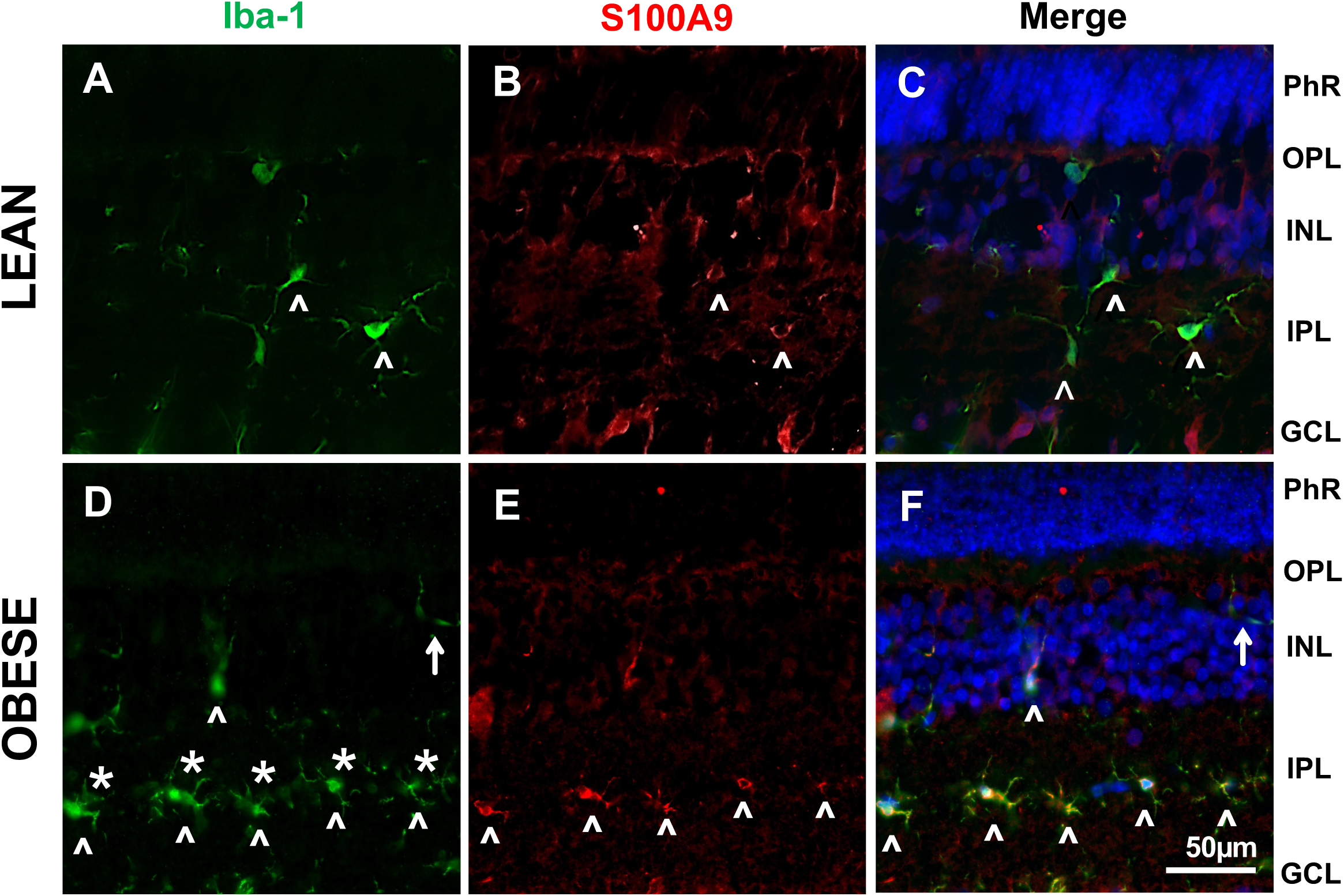
S100A9 colocalized with Iba+ microglial cells. Double immunostaining of (**A, D**) Iba-1 for microglia and (**B, E**) S100A9 found the protein (^) to be present in the cytoplasm of microglial cells, which was amoeboid-like (**\***) in the obese Ossabaw pig retina. S100A9 was also found upregulated (↑) in the microglial cells present in the OPL of the obese pigs (**F**).

### pMicroglia produced S100A9 in vitro

pMicroglia exhibit ramified morphology *in vitro*, consistent with their structure in naïve pig retina. LPS is adopted as an activating stimulus, transforming the pMicroglia into amoeboid structures after 24h (Fig 3A-D). When evaluated for S100A9 protein, activated pMicroglia increased S100A9 transcript expression in a dose-dependent manner (Fig. 3E), which was translated to monomeric and dimeric proteins intracellularly (Fig. 3F). The DAMPs molecule was also secreted into the culture media at a 4-fold increase after LPS stimulus (Fig. 3G). These results supported the *in vivo* data that pig microglial cells could be one of the major sources of S100A9 in the diabetic retina.

**Figure 3.**
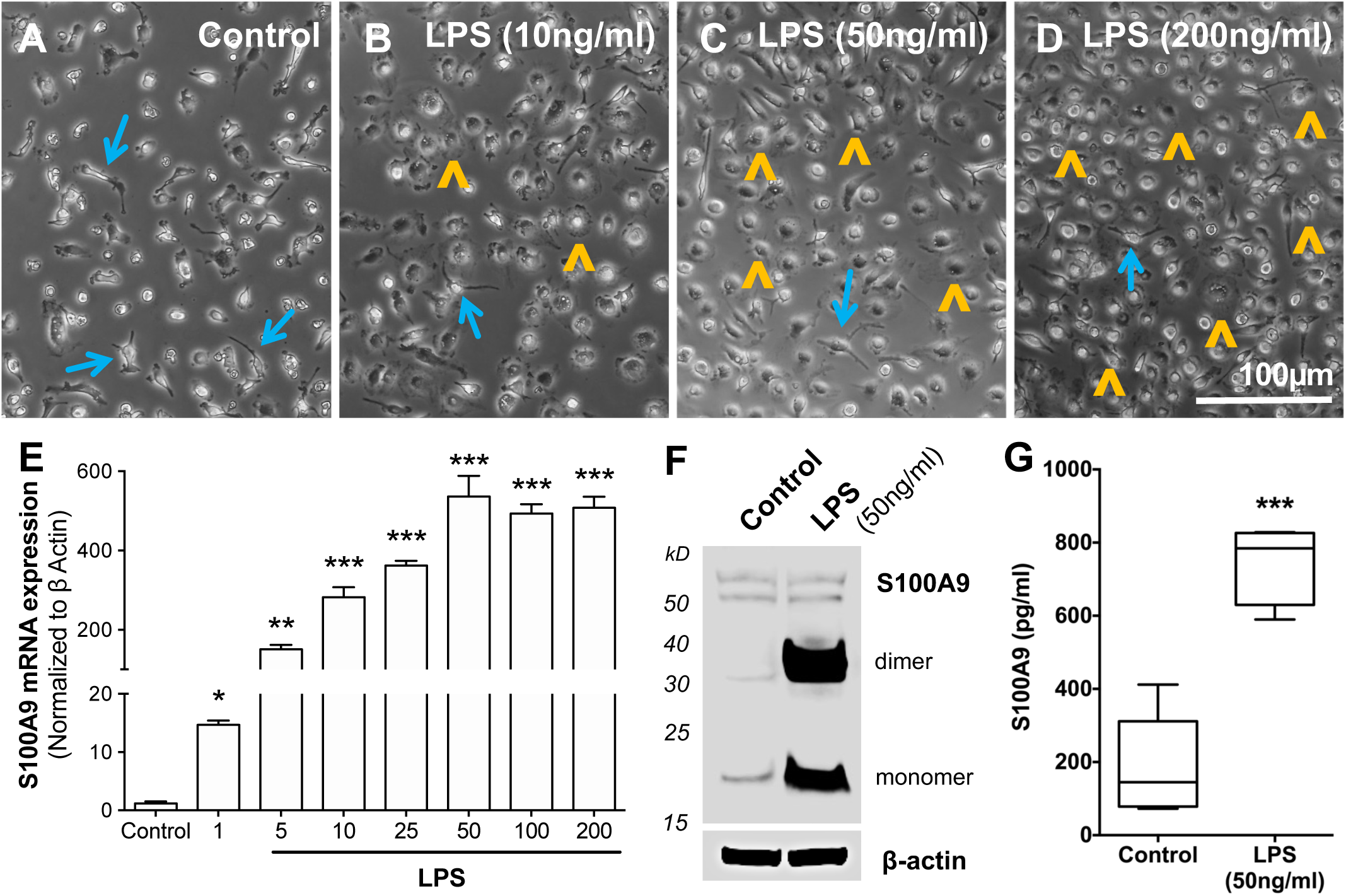
S100A9 production in pMicroglia. pMicroglia treated with LPS for 24 hours showed the morphological transformation from (**A-D**) the ramified state (↑) to amoeboid (^) phenotype from 10ng/ml onwards. (**E**) S100A9 transcripts were upregulated in a dose-dependent manner, reflected by a significant increase in (**F**) intracellular and (**G**) secreted S100A9 protein in the culture media. *, P<0.05; **, P<0.01; ***, P<0.001.

### S100A9 induced pro-inflammatory response from pMicroglia

Since S100A9 accumulation has been reported to cause sterile inflammation in other diseases^50^ we wanted to understand how the microglial cells respond to excess DAMPs protein in the tissue. Therefore, pMicroglia were treated with recombinant human S100A9 (R&D Systems) for 24 hours. Cells showed amoeboid morphology as seen with LPS, indicating a microglial shift to the activated state (Fig. 4A). Likewise, S100A9 treatment resulted in a dose-dependent increase of IL-1β transcripts (Fig. 4B) with an ED_50_ of approximately 89.14nM (Fig. 4C). The pro-inflammatory cascade was also seen by a significant production of pro-IL-1β, which was cleaved to form the active IL-1β only in the S100A9 treated cells (Fig. 4D). S100A9 treated pMicroglia also showed de novo synthesis of S100A9 transcripts (Suppl Fig. 2), as well as NLRP3 and TLR4 mRNA (Fig. 4E). S100A9 was reported to signal through TLR4 binding, and NLRP3 inflammasome was previously shown to process IL-1β cleavage. NLRP3 protein was elevated after S100A9 treatment, but TLR4 expression was not observed to increase at the protein levels (Fig. 4F).

**Figure 4.**
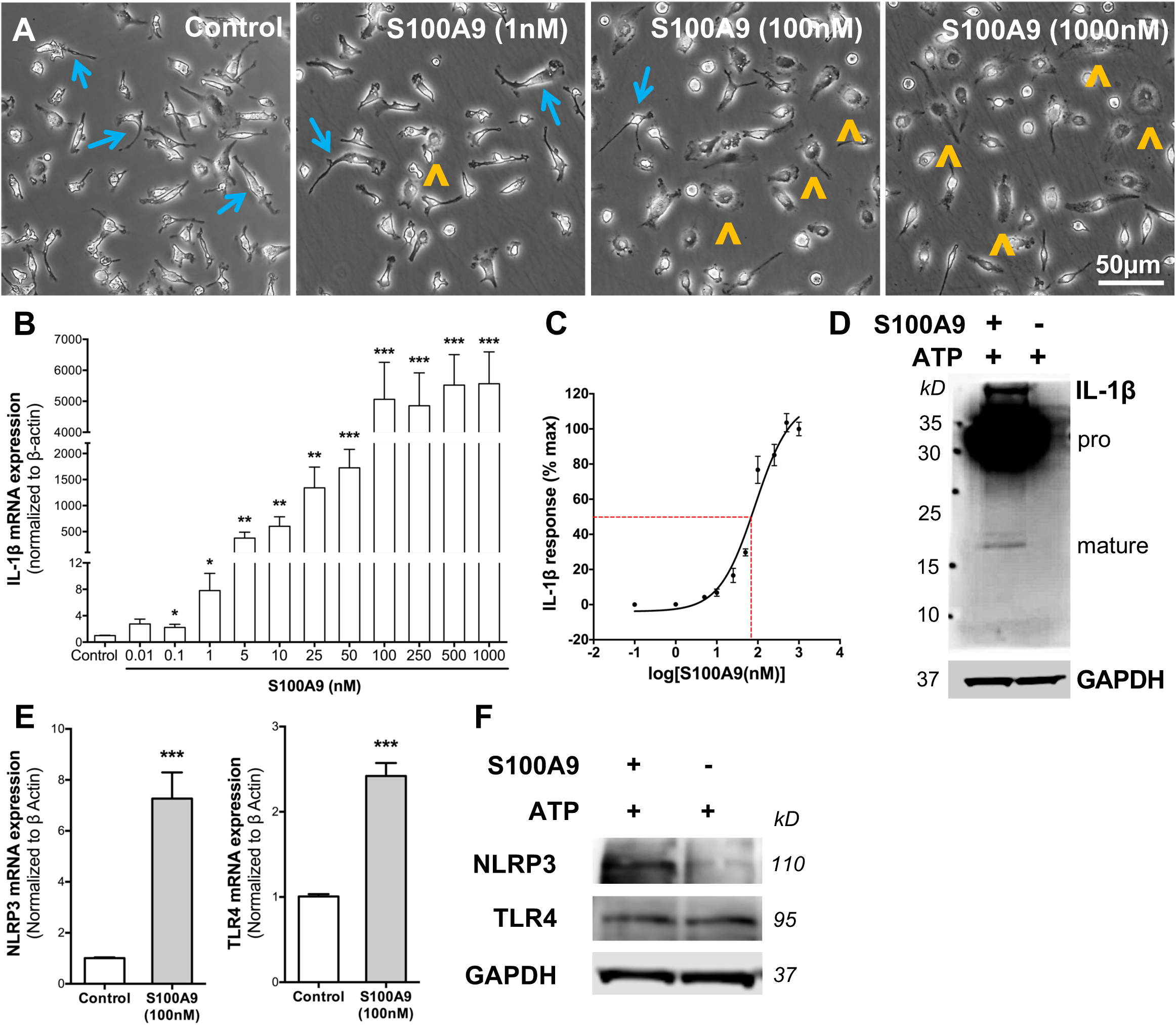
S100A9 induced pro-inflammatory response in pMicroglia. pMicroglia seeded at 5×10^4^ cells/cm^2^ density were treated with exogenous recombinant S100A9 (0.01nM to 1000nM) in DMEM-HG on day 2 and collected after 24 hours for qPCR and western blot. (**A**) Representative phase contrast images of pMicroglia show morphological change from ramified state (↑) to amoeboid (^) phenotype treated with S100A9 at 100nM. (**B**) Real-time PCR showed IL-1β transcript increase in a dose dependent manner, with (**C**) estimated ED_50_ of 89.14nM. (**D**) Western blot also showed cleaved IL-1β protein in the S100A9 (100nM) treated group. (**E**) NLRP3 and TLR4 was also increased at the RNA level, however only (**F**) NLRP3 protein was seen to be increased in western blot. *, P<0.05; **, P<0.01; ***, P<0.001.

### S100A9 functions via NLRP3 inflammasome in microglial cells

Inhibition studies were performed to examine the mechanism of action further. TAK242 is a specific inhibitor of TLR4 that binds to the intracellular domain of the protein to disrupt the interaction of the receptor with adaptor proteins^51^. TAK242 pre-treatment prior to S100A9 stimulus reduced the expression of glycosylated-TLR4, seen as the 120kD band in immunoblots (Fig. 5A). Glycosylation of TLR4 had been reported to be essential for cell surface delivery of the receptor, and proper folding into its active conformation^52^. Decreased active TLR4 could indicate attenuation of S100A9 activity, which was reflected by reduction in IL-1β secretion (Fig. 5B). Since IL-1β is known to be processed by the NLRP3 inflammasome, we also used MCC950, a specific inhibitor of NLRP3 inflammasome to examine the contribution of the inflammasome complex to IL-1β production following S100A9 induction. MCC950 binds to the NACHT domain of the protein, thereby preventing it from assuming an open conformation ready for ATP hydrolysis, a crucial step in NLRP3 activation^53^. Recombinant ATP (5mM) was therefore given to cells 15 minutes prior to collection. S100A9 treatment resulted in increased NLRP3 but decreased ASC expression (Fig. 5C), but did not hamper IL-1β production (Fig. 5D). However, MCC950 pre-treatment reduced NLRP3 and ASC expression (Fig. 5C), which translated to a significant attenuation of secreted IL-1β (Fig. 5D). Taken together, inhibitor studies support the hypothesis that S100A9 binds to TLR4 in microglial cells and activates NLRP3 for production of IL-1β.

**Figure 5.**
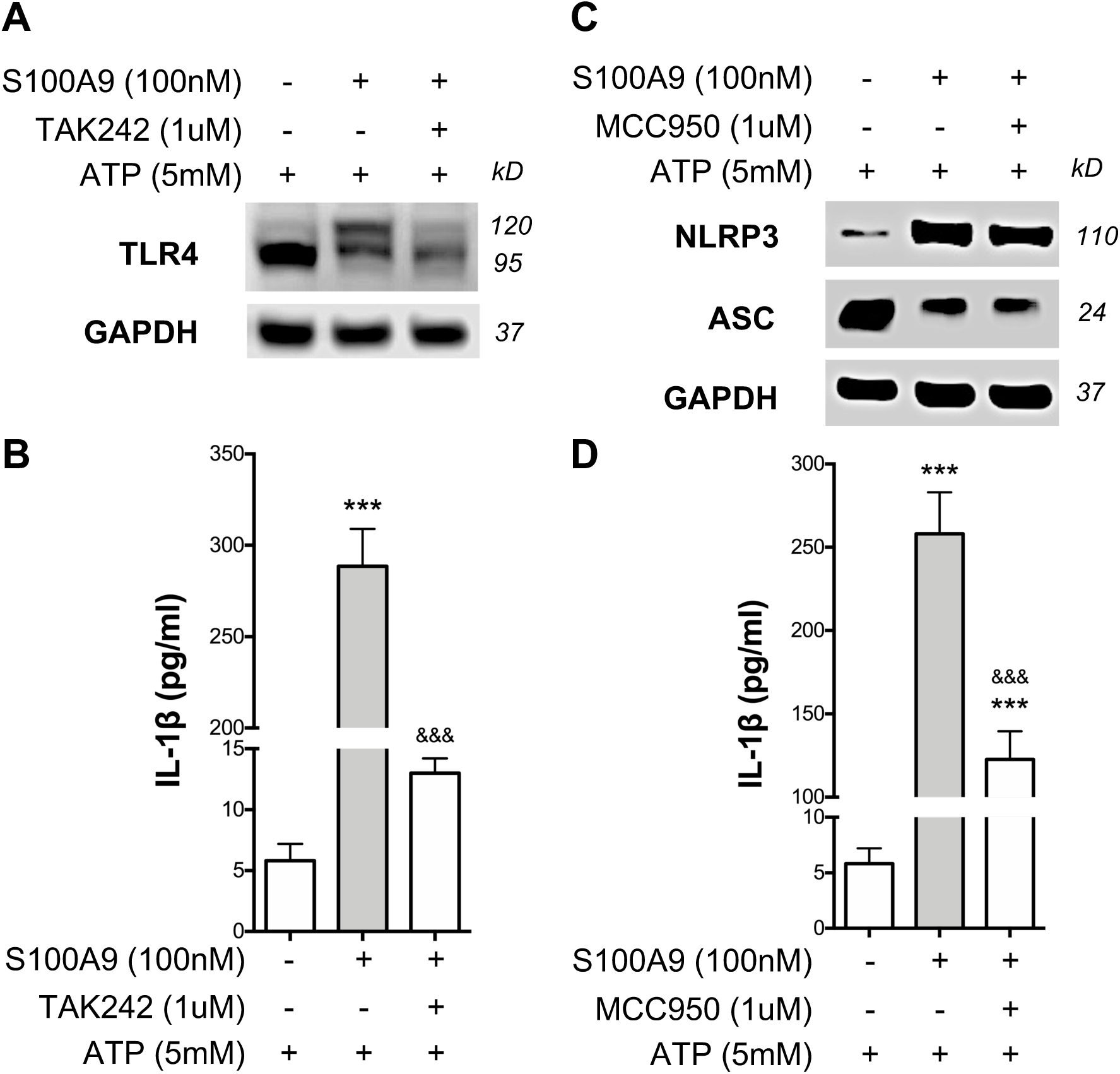
S100A9 functions via NLRP3 inflammasome. Microglial cells were pre-treated with TLR4 inhibitor – TAK242 (1μM), or NLRP3 inhibitor – MCC950 (1μM) for 30 minutes prior to addition of recombinant S100A9 protein. Cells were collected after 24 hours for western blot, and cell culture supernatant was evaluated for IL-1β secretion by ELISA. (**A**) S100A9 treatment resulted in glycosylation of TLR4 receptor, seen as the 120kD band on Western blot, which was attenuated with TAK242. (**B**) Microglial cells secreted significantly high levels of IL-1β upon S100A9 stimulus but was significantly reduced after partial inhibition of the TLR4 receptor by TAK242. Similarly, (**C**) MCC950 treatment reduced the activation of NLRP3 inflammasome, as seen in reduced NLRP3 and ASC, and **D**) decreased IL-1β secretion after 24 hours. The experiment was performed three times with n=3. Data are represented as mean±SEM. ***, P<0.001 compared to control; ^&&&^, P<0.001 compared to S100A9.

### Paquinimod attenuates S100A9 activity in microglial cells

Paquinimod (ABR-215757), is a quinoline-3-carboxamide (Q-compound), which interacts with S100A9 and inhibits its binding to TLR4^54^. The family of Q-compounds act as immunomodulatory compounds due to the regulation of cellular immune and cytokine responses without suppression of adaptive immunity^55^. Paquinimod was shown to be safe and tolerable in clinical studies of systemic lupus erythematosus^56^ and was recently reported to prevent diabetes progression in the non-obese diabetic (NOD) mouse model of T1DM^57^. Despite its known benefits in modulating the innate immune cells, the effect of Paquinimod on microglial cells have not yet been evaluated.

We first established the safety of Paquinimod by treating microglial cells with increasing doses of the drug (Fig 6A-D). Paquinimod did not cause a reduction in cell viability in concentrations below 300μM, as seen with PrestoBlue assay (Fig. 6E). Effectiveness of the Q-compound was next evaluated in the morphological and biochemical analysis of microglial cells with Paquinimod prior to S100A9 stimulus. While S100A9 induced morphological changes in the microglial cells by furling the cell edges and flattening of cell bodies (Fig. 6B), Paquinimod prevented the transformation of microglia (Fig. 6C, D). The drug also reduced IL-1β (Fig. 6F) and S100A9 (Fig. 6G) transcripts in a dose-dependent manner. In addition, paquinimod decreased NLRP3 components – NLRP3, ASC, and Caspase-1 proteins (Fig. 7A). This suggests that Paquinimod is an effective inhibitor that prevents binding of S100A9 in the microglial cells, which led to an overall attenuation of IL-1β secretion (Fig. 7B). Altogether, Paquinimod is safe for application to microglial cells, and was effective to limit S100A9-induced inflammation, via downregulation of NLRP3 inflammasome activity.

**Figure 6.**
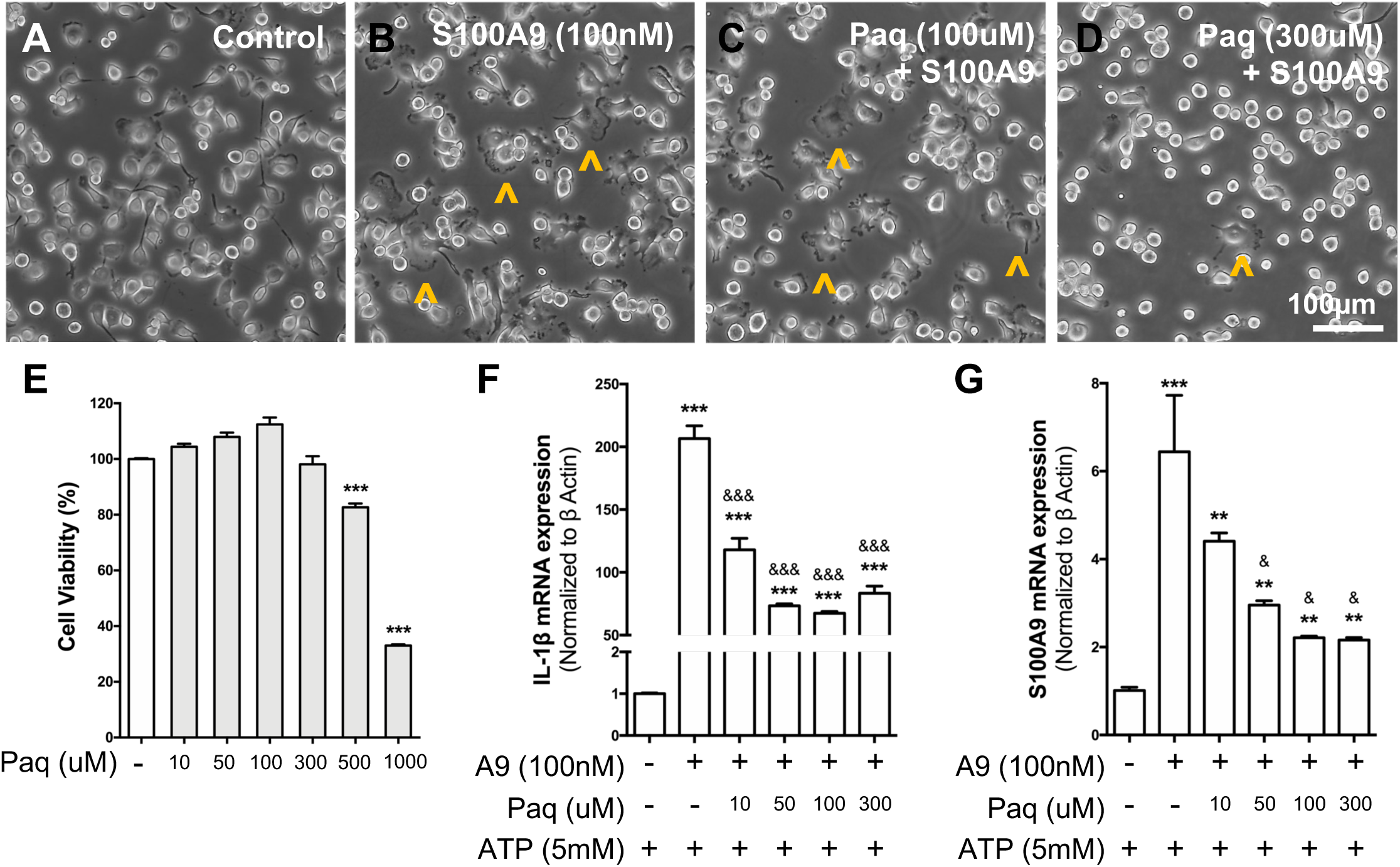
Paquinimod is safe in Microglial cells and reduces IL-1β and S100A9 production in a dose-dependent manner. Paquinimod (Paq), a quinoline-3-carboxamide (Q compound) that inhibits S100A9 (A9) binding to TLR4, was treated 30 minutes prior to the addition of S100A9 in microglial cells. (**A-D**) Representative phase contrast images of (**B**) S100A9 treated cells after 24 hours showed furling of the cell boundaries and flattening of cell body (**^**), indicative of activated morphology. (**C,D**) Paquinimod (100μM or 300μM) attenuated the morphological changes without cell death. (**E**) PrestoBlue cell viability assay indicates toxicity of Paquinimod above 500uM after 24 hours of treatment. Transcript expression of (**F**) IL-1β and (**G**) S100A9 was attenuated in a dose-dependent manner with Paquinimod pre-treatment. *, P<0.05; **, P<0.01; ***, P<0.001 against control. ^&^, P<0.05; ^&&^, P<0.01; ^&&&^, P<0.001 against S100A9 (100nM).

**Figure 7.**
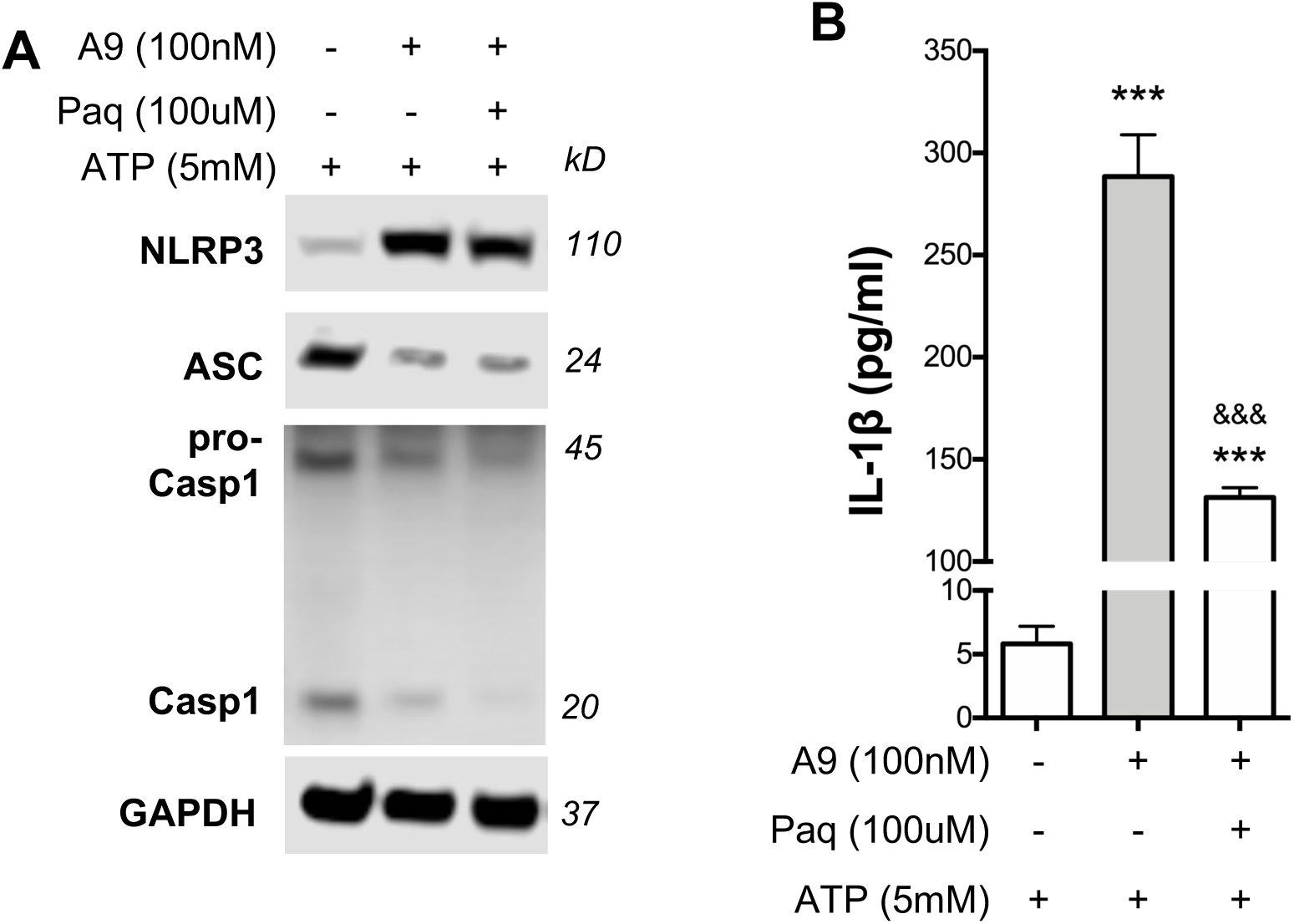
Paquinimod exert inhibitory functions on S100A9 via NLRP3 inflammasome. Microglial cells pre-treated with Paquinimod (Paq) for 30 minutes prior to S100A9 (A9) stimulus were collected after 24 hours for Western blot and ELISA. (**A**) Paquinimod reduced NLRP3 inflammasome components, including NLRP3, ASC, and Casp1 proteins. (**B**) IL-1β secretion was also significantly reduced with Paq treatment. ***, P<0.001 compared to control; ^&&&^, P<0.001 compared to S100A9.

## Discussion

DAMPs-driven sterile inflammation in diabetic retinopathy is a novel mechanism that is not yet fully understood^58^ (Mahaling et al. 2023). Inflammation prepares the body for defense against insult and injury, but uncontrolled prolonged endogenous danger signals can exacerbate the repair process, causing deleterious effects^59^. S100A9 is the most abundant DAMPs in many inflammatory disorders, including rheumatic diseases^26^ and inflammatory bowel disease^25^. It was previously found in the circulation of T2DM patients with DR^30^ and subsequently shown in the epiretinal membranes of human PDR patients^19^. The present study showed the presence of S100A9 in the retinal microglial cells of the obese diabetic Ossabaw pig, indicating its potential to instigate sterile inflammation in the retina. We further demonstrated S100A9 production from cultured retinal microglial cells and inhibited its pro-inflammatory action with Paquinimod, an immunomodulatory Q compound.

Sterile inflammation in the retina during DR is recognized by elevated expressions of pro-inflammatory mediators such as IL-1β, IL-6, IL-8^4^, and DAMPs^18,36,58^. In Ossabaw pigs fed a Western diet for 2.5 months, we found that S100A9 was expressed by Iba-1 stained microglial cells and supported by the brain microglia transcriptomics^35^. Notably, the expression of S100A9 was elevated in microglia residing in the OPL of the retina, indicative of microglial response to early retinopathy. While multiple modes, including extracellular matrix degradation, cell death, and autophagy have been proposed for DAMPs production in retinal diseases^60^, the mechanism of S100A9 secretion in the retina during DR remained to be studied. To further delineate the mechanism of S100A9 action during DR, we stimulated primary porcine retinal microglial cells with LPS, resulting in elevated production of S100A9, supporting their use as an *in vitro* model for studying the role of microglia in retinal diseases.

Pro-inflammatory functions of S100A9 have been documented in several systemic diseases. Not only does it promote the release of IL-1β from macrophages^61^, but it also serves as a chemotactic factor for macrophage and neutrophil infiltration^62^, and increased permeability in endothelial cells^63^. In the present study, treatment of microglial cells with S100A9 induced an inflammatory response by a morphological shift from ramified to amoeboid state and heightened the production of IL-1β, which has been implicated in DR pathogenesis. S100A9 is known to bind TLR4 receptors and initiate NF-kB-mediated transcription of pro-inflammatory cytokines^64^. In chronic hyperglycemia, microglia were continually activated, causing inflammatory milieu^6^ throughout the DR progression as reported in PDR patients^30^. Thus, persistent S100A9 production could result in constant NLRP3 activation, thereby causing sterile inflammation in the retina.

Attenuating DAMPs is beneficial in ameliorating inflammatory diseases. For instance, HMGB1 inhibition protected the retina against vascular degeneration in streptozotocin (STZ)-induced diabetic mice^65^. HSP90 inhibition also suppressed neovascularization in diabetic mice, 1-week after STZ induction^66^. Overall blockade of TLR4 activation in diabetic rats with TAK242 administration reduced leukocyte infiltration and attenuated blood-retinal breakdown via downregulation of the MyD88 pathway^67^. In this study, blocking TLR4 using TAK242 partially prevented S100A9-induced IL-1β production. The increase in TLR4 following S100A9 treatment was also inhibited with TAK242. These results are consistent with the hypothesis that S100A9 binds to TLR4 in the microglial cells. Meanwhile, since NLRP3 inflammasome was shown to contribute to DR progression^17,36^, MCC950 was used to evaluate the downstream involvement of the inflammasome complex in response to microglial activation. MCC950 binds to NLRP3 and prevents ATP hydrolysis, which is required for its activation^53^. MCC950 treatment decreased the S100A9-induced production of NLRP3 protein and attenuated IL-1β release, indicating the direct role of S100A9 in IL-1β production via NLRP3 inflammasome in the microglial cells.

Since targeting TLR4 affects other TLR4-dependent defense mechanisms^68^, we used an S100A9-specific inhibitor, Paquinimod (ABR-215757), in the microglial cells. A previous Phase I study in patients with systemic lupus erythematosus (SLE) found Paquinimod to be safe and effective, resulting in reduced serum interferon alpha (IFN) score after 12 weeks of oral administration^56^. In another study, Paquinimod intervened in leukocyte recruitment in an experimental peritonitis model without affecting the dynamics of the myeloid population^55^. Interestingly, Paquinimod also reduced the recruitment of inflammatory monocytes from the blood vessels in the same study. In the non-obese diabetic (NOD) mice model, Paquinimod-administration reduced diabetes development from 73% to 30%^57^. *In vitro*, synovium from humans with end-stage osteoarthritis (OA) treated with Paquinimod in the presence of S100A9 resulted in reduced expression of IL-6 and IL-8^69^. Mechanistically, experiments suggest Paquinimod to compete for S100A9 binding to TLR4^54^. However, the exact binding site of the drug to the S100A9 complex is not yet elucidated^68^. We observed that pre-treatment of microglial cells with Paquinimod prior to S100A9 treatment reduced TLR4, inhibited NLRP3 inflammasome complex, and attenuated IL-1β production (Fig. 8).

**Figure 8.**
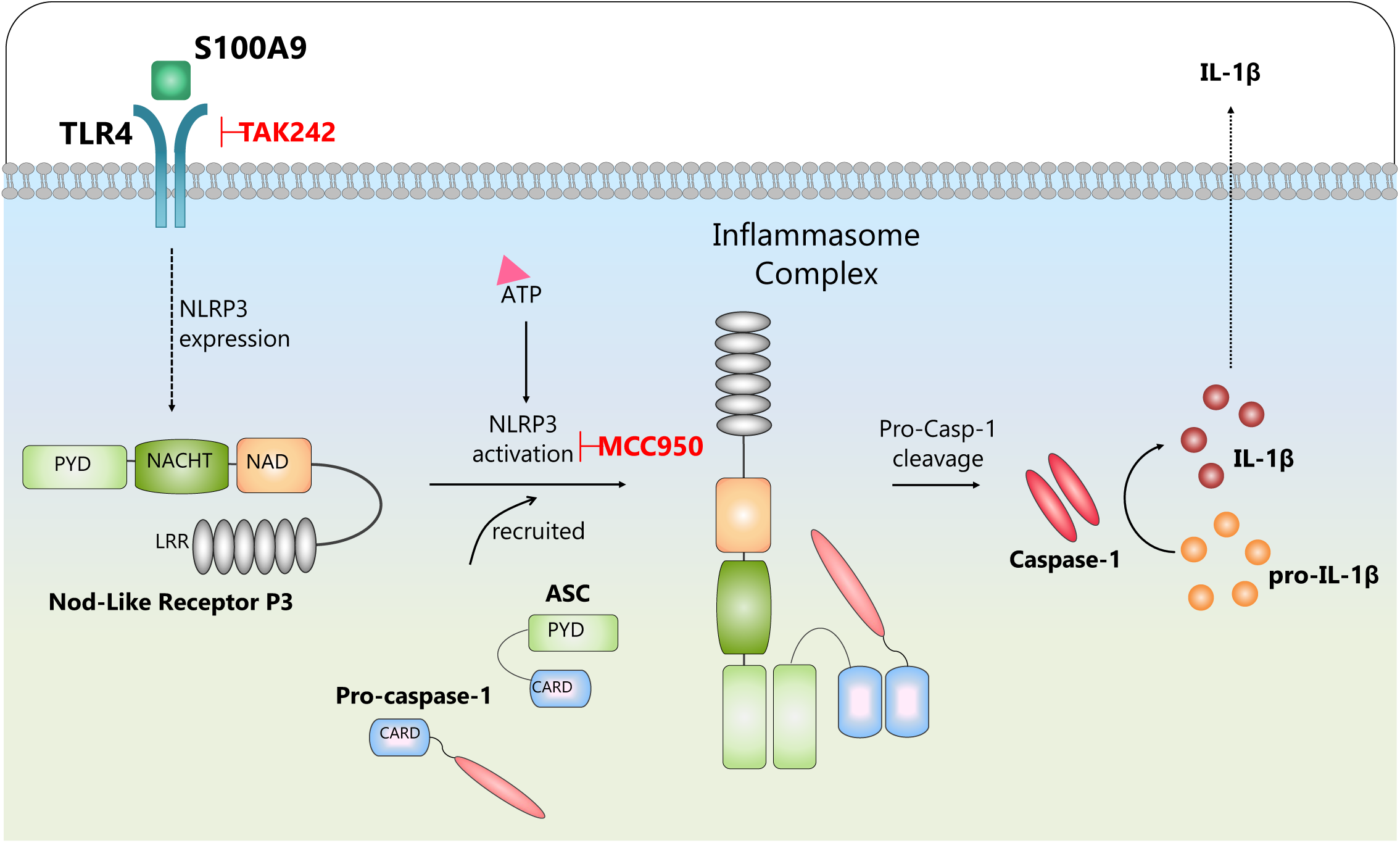
Schematic depicting the mechanism of action of S100A9 signaling pathway in the Pig Retinal Microglial cells.

In summary, we found that S100A9 is upregulated in the Ossabaw pig model of early DR produced by activated retinal microglial cells. S100A9 induce IL-1β release, and inflammatory response through the activation of NLRP3 inflammasome was attenuated using an S100A9-specific inhibitor, Paquinimod. Thus, this study demonstrated S100A9 as a possible target for intervention in the progression of DR.

## Acknowledgments

Supported by funding from National Institutes of Health (NIH)/National Eye Institute (NEI) grant R01 EY029795 to S.S.C.

## Disclosures

The authors declare no competing interests.

## Notes

### Competing Interest Statement

The authors have declared no competing interest.

