## Supplemental File 1 for "Retinal microglia-derived S100A9 incite NLRP3 inflammasome in a Western diet fed Ossabaw pig retina"

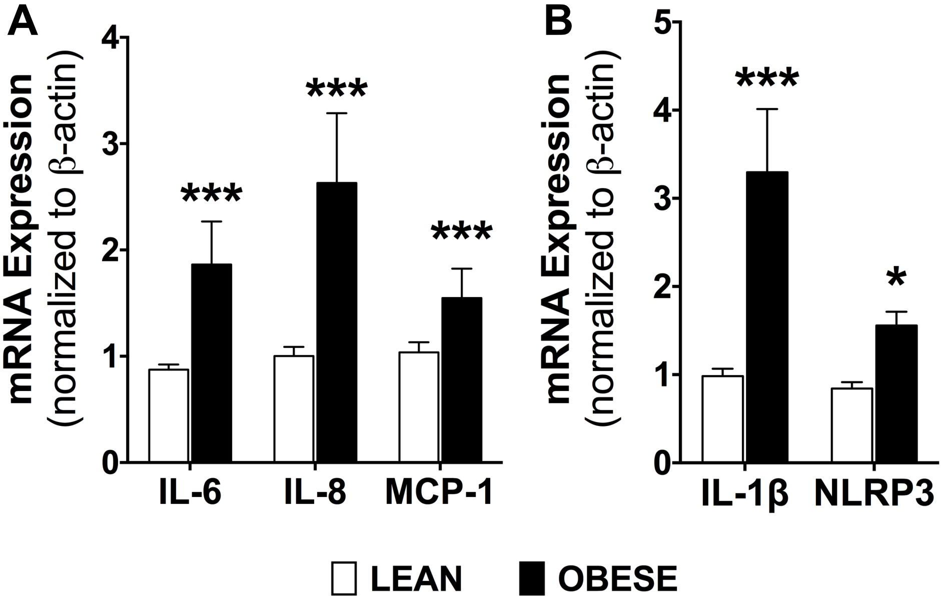


**Supplementary Figure 1. Ossabaw pigs fed a western diet showed increased levels of inflammatory factors and mediators in retina.**
(**A**) Real-time PCR on obese Ossabaw pig retina showed increased production of IL-6, IL-8 and MCP-1 transcripts. (**B**) IL-1β and NLRP3 were similarly elevated. n = 3 eyes per group. *, $P<0.05$; ***, $P<0.001$.
