## Supplemental File 2 for "Retinal microglia-derived S100A9 incite NLRP3 inflammasome in a Western diet fed Ossabaw pig retina"

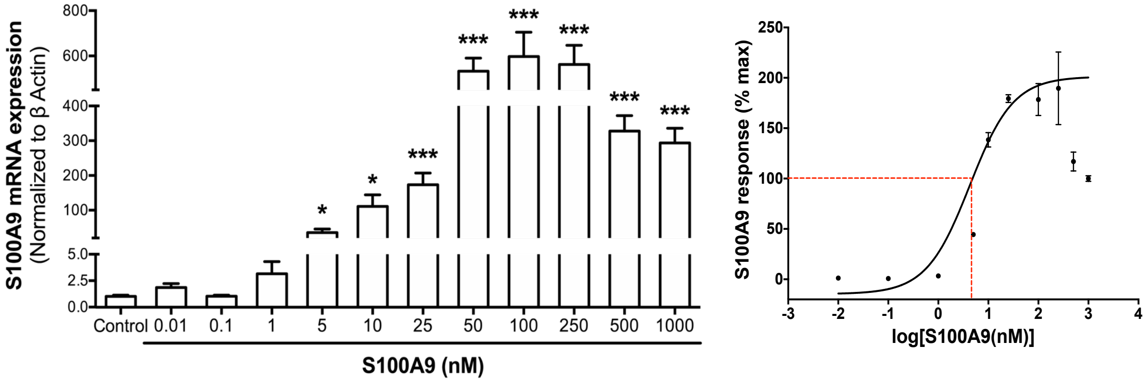


**Supplementary Figure 2. Autocrine expression of S100A9 transcripts.** pMicroglia treated with increasing doses of S100A9 for 24 hours showed dose-dependent increased expression of S100A9 transcripts. *, P<0.05; ***, P<0.001.
